# Multimodal induction of fulminant HLH by IL-18 includes virus-specific NK immunodeficiency

**DOI:** 10.1101/2025.07.04.663237

**Authors:** Jemy Varghese, Emily Landy, Leonardo Huang, Anastasia Frank-Kamenetskii, Hannah Klinghoffer, Jeremy Morrissette, Vinh Dang, Stephen Carro, Laurence Eisenlohr, Scott Canna

## Abstract

Macrophage Activation Syndrome (MAS) is a cytokine storm syndrome associated with Still’s Disease, XIAP deficiency, and elevation of both total and free IL-18. Modeling excess IL-18 using *Il18tg* mice, we found mild NK-cytopenia and cytotoxic T lymphocyte (CTL) activation in resting mice reminiscent of Still’s patients. Infection with Lymphocytic Choriomeningitis Virus (LCMV) triggered MAS via and IFNγ, despite normal viral clearance. *Il18tg* NK cell transcriptomes showed replicative activity, but few changes in canonical NK function or maturation pathways. LCMV clearance is NK-independent, so we challenged *Il18tg* mice with mousepox in which NK cells are critical early orchestrators of clearance. *Il18tg* mice’s NK cells failed to activate or expand, but mousepox further activated their CTL and early viral control was normal. *Il18tg* mice soon developed “MAS” including hepatosplenic necrosis, but (contrasting with LCMV) they showed poor virus-specific CTL expansion and viral clearance. Though more normal at rest, *Il18bp^KO^* mice’s NK cells were similarly inert upon mousepox infection, and the mice succumbed to viremic MAS like *Il18tg*. Rescue of *Il18tg* mice, and mousepox-specific CTL responses, by NK cell transfer required in vitro NK pre-activation. Thus, IL-18 can induce both hyperinflammation (CTL *hyper*activation) and immunodeficiency (NK cell *hypo*activation) depending on the nature of the infectious trigger.

## Introduction

Hemophagocytic Lymphohistiocytosis (HLH) is a cytokine storm syndrome of systemic inflammation, coagulopathy, cytopenias, hepatitis, and hyperferritinemia. Mechanistically, HLH is canonically associated with biallelic defects that cripple granule-mediated cytotoxicity (GMC), as seen in Familial HLH (FHL). However, the great majority of patients meeting established clinical criteria have intact GMC. These patients’ secondary HLH (sHLH) arises in infectious, malignant, cancer immunotherapy, and rheumatic contexts^1^. The latter, better known as Macrophage Activation Syndrome (MAS), is largely described in Still’s Disease and Systemic Juvenile Idiopathic Arthritis where it is strongly linked to the cytokine IL-18^2^.

Infection-associated HLH (IA-HLH) occurs in patients with otherwise normal-appearing immune systems. Nearly any pathogen can serve as a trigger, but primary EBV infection is by far the best-described, followed by other DNA viruses and (largely intracellular) microbes^1^. The distinction between IA-HLH and hyperinflammatory sepsis is unclear^3^. Infection is a common HLH trigger regardless of context. Children with FHL typically appear healthy at birth.

Symptoms usually present in the first year of life, presumably triggered by infection (though often none is identified), but reports of later presentations are plentiful. EBV is also a classic trigger for primary HLH in genetically-susceptible individuals (e.g. *SH2D1A*, *XIAP*, *CD27*, …)^4–7^. In MAS, at least 34% of episodes involve an infectious trigger, and again EBV is reported most commonly^8^.

Seeking pathoetiologic overlap between MAS and FHL, studies in MAS have noted enrichment for rare heterozygous variants in GMC genes associated with FHL^9–10^. MAS patients also showed NK cytopenia and impaired NK cell function^12–16^. However, more recent work has emphasized a unique role for IL-18 in MAS. IL-18 is an inflammasome-activated cytokine best known for acting on NK cells, activated/memory T-cells, and Chimeric Antigen Receptor (CAR)

T-cells to induce Interferon-γ (IFNg) and promote GMC^2,17^. Profoundly elevated levels of IL-18 often exceed neutralization by IL-18 binding protein (IL-18BP) and serve as a uniquely specific diagnostic biomarker of MAS-prone disorders, so-called “IL-18opathies”^2,18–21^. The discovery of several monogenic autoinflammatory IL-18opathies^2,22–26^, and a small but growing experience with IL-18 blockade^27–29^, help place excess IL-18 upstream of MAS.

Animal models of HLH/MAS have been essential to our understanding of their complex immunobiology. Lymphocytic Choriomeningitis Virus (LCMV) is a common mouse pathogen cleared readily in wild-type (wt) mice. As in human FHL type 2^30^, perforin-deficient mice appear normal at rest, but LCMV infection leads to persistent viremia and drives murine HLH via unopposed antigen presentation, CD8 T-cell hyperactivation, and IFNg overproduction^31,32^.

Similarly, mice with transgenic IL-18 expression (*Il18tg*) also develop normally and LCMV infection also drives CD8 T-cell hyperactivation, IFNg overproduction, and HLH/MAS. Unlike *Prf1^KO^* mice, however, *Il18tg* mice clear LCMV normally^33–35^ and their T-cell contraction and memory formation are ultimately normal^33^. In mice and in vitro, IL-18 synergizes with GMC impairment and preferentially promotes CD8 T-cell responses^2,33–35^. Given NK cells’ constitutive expression of the IL-18 receptor and the NK cell abnormalities observed in Still’s patients, we sought to evaluate NK cell characteristics in mice with excess IL-18 at rest and upon systemic NK challenge.

## Materials and Methods

### Mice, Infections, & NK Depletion

Adult mice aged 8-13 weeks were maintained, bred, and handled in specific pathogen free conditions under animal protocols approved by The University of Pittsburgh or The Children’s Hospital of Philadelphia. Male and female mice were distributed equally across groups and analyzed together. Wild-type (WT) and *Il18^KO^* mice were obtained from Jackson Laboratories. *Il18tg* mice were a gift from Tomoaki Hoshino (Kurume University)^36^, *Il18bp^KO^* mice (Il18bp^tm1.1(KOMP)Vlcg^) were obtained from the knockout mouse project. Mice were genotyped by PCR using appropriate primers (**Supplemental Table 1**).

Mousepox ectromelia virus (ECTV, Moscow strain, 3000 pfu via hind footpad) infections were performed as in previous publications^33,37^. To deplete NK cells, mice were injected IP with 200ug of anti-NK1.1 (BioXCell, BE0036; clone PK136) or rat IgG2a isotype control (BioXCell, BE0090; clone LTF-2) −1 and +2 days after infection with ECTV.

### Flow cytometry

Organs were weighed immediately ex vivo. Splenocytes were digested in media (DMEM+2%FBS) containing collagenase D (0.135U/ml) and DNAse (100 μg/ml) for 25-30 minutes, followed by mechanical disruption and osmotic (ACK) lysis. Livers were perfused in situ, collected into media, homogenized using gentleMACS tissue dissociator (Miltenyi), and digested in collagenase D and DNAse as above. Bone marrow cells were isolated from tibias and/or femurs through hydrostatic extrusion using media, followed by mechanical disruption and RBC lysis. Whole blood was collected into EDTA tubes, plasma isolated by centrifugation, and peripheral blood mononuclear cells (PBMC) isolated using Ficoll (GE Healthcare). Live cells were counted by Acridine Orange/Propidium Iodide on Nexcelom Cellometer K2.

Single cell suspensions were stained in HBSS for 30 minutes using antibodies/reagents listed in the **Supplemental Methods**. Cells were acquired on a 4-laser Beckman Coulter CytoFLEX S, 6-laser Beckman Coulter CytoFLEX LX, or five-laser Cytek Aurora system and analyzed using FlowJo version 10.10.0 (Treestar). UMAP was generated using UMAP 4.1.1 plugin and R version 4.4.1. UMAP was run with setting as: number of nearest neighbors: 15; minimum distance: 0.5, distance metric: euclidean; supervised weighting 0.1. Parameter used for analysis include B8R tetramer, FasL, CD44, TIGIT, Lag-3, CD11b, IL-18Rα, CD39, and PD-1.

### In vivo killing assays

These were performed at day 7-8 post infection as in previous publications^33,38^ using congenically-marked splenocytes loaded with ECTV B8R^20–27^ peptide and labeled with Tag-it Violet (Biolegend), co-transferred 1:1 with CellTrace™ CFSE (Invitrogen) labeled control splenocytes (total 8-10×10^6^ cells/mouse). Transferred cells were quantified by flow cytometry in spleens of recipient mice 4 hours post-transfer.

### Virus Quantitation

Total RNA from spleen and liver was obtained using Trizol (Invitrogen) and Chloroform (Fisher Scientific) per manufacturer’s instructions. First-strand cDNA was synthesized with qScript™ cDNA Synthesis Kits, Quantabio. For 18srRNA and Evm003 RT-qPCR was performed using Quantabio PerfeCTa® SYBR® Green SuperMix, Quantabio. with the following primers: Evm003: F: TCT GTC CTT TAA CAG CAT AGA TGT and R: TGT TAA CTC GGA AGT TGA TAT GGT A, 18srRNA: Applied Biosystems Taqman reagent mix (Hs99999901_s1). RT-qPCR was done using (QuantStudio™ 5 System) and 2^deta CT was calculated (ECTV-18s rRNA). ECTV with replicative potential was assessed by plaque assay on BS-C-1 cells (as in *Peauroi et al*^37^) from the same organs assessed by qPCR (Supplemental Figure 3).

### Histology

Tissues were fixed in 10% buffered formalin (Fisher SF98-4) for 24 hours prior to processing, paraffin embedding, and cutting into 5um sections. Slides were stained with hematoxylin and eosin or Trichrome Stain LG Solution (Sigma Aldrich HT10316). Slides were scored for liver damage by two independent, blinded reviewers using Leica DM4000B upright light microscope and Leica DFC7000 T camera. Liver damage was estimated at 400x magnification by the following criteria: (1) 0-25% damage; (2) 25-50% damage; (3)50-75% damage; (4) 75-100% damage. Additionally, necrosis score (0-2) was added to the score to better represent the tissue damage.

### RNA-seq

Splenic NK (Live, TCRβ-, B220-, NK1.1+, NKp46+), liver NK (Live, TCRβ-, B220-, NK1.1+, NKp46+, TRAIL-), and Liver ILC1 cells (Live, TCRβ-, B220-, NK1.1+, NKp46+, TRAIL+) were purified by double-sorting and ∼1,000 cells were sorted directly into Smart-Seq V4 ultra-low-input RNA sequencing preparation kit lysis buffer. mRNA purification and fragmentation, cDNA synthesis, and target amplification were performed per manufacturer’s instructions (Takara). Pooled cDNA libraries were sequenced on the Illumina NextSeq500, mapped to Mus_musculus_ensembl_v80 reference sequence and gene track Mus_musculus_ensembl_v86, and quantified using CLC Genomics Workbench software (Version 22, Qiagen). Differentially expressed genes were defined as having a maximum group mean transcripts per million (TPM) ≥ 5, absolute fold change ≥2, and false discovery rate ≤0.01. Fold changes were not calculated directly from transcripts per million values but from the generalized linear model, which corrects for differences in library size between the samples and the effects of confounding factors. Canonical NK cell transcripts were identified from MsigDB pathways (KEGG_NATURAL_KILLER_CELL_MEDIATED_CYTOTOXICITY, BIOCARTA_NKCELLS_PATHWAY) and Bezman et al.^39^. See **Supplemental Methods** for details on generation of “NK IL-18 Response” and “Immature NK/ILC” custom gene sets.

### Gene Set Enrichment Analysis (GSEA)

GSEA comparing *Il18tg* vs WT for each cell type was performed using GSEAv4.2.3 (after removing genes with group mean TPM<5) using the following parameters: Gene sets=MSigDBC7.all.V7.1 (Immunologic Signatures), C2 (Canonical Pathways), or custom gene sets as above; number of permutations=1000; permutation type=gene_set. Heat maps were generated using Morpheus (Broad Institute).

### NK culture and Expansion

Splenic NK cells were purified using MojoSort negative selection kit (Biolegend). To assess survival ex vivo, cells were cultured in complete medium (**Supplemental Methods**) for 5 hours and subsequently assessed for NK cell identity and viability by flow cytometry. For ex vivo activation/expansion, WT splenic NK cells were purified as above and cultured in complete medium supplemented with murine IL-2 (1000U/mL, PeproTech). Cells were split and media replaced every 3 days for 2 weeks, washed twice in PBS and injected IV on the day prior to infection (5×10^6^/mouse).

### Statistical analyses

Frequentist statistical analyses were performed in GraphPad Prism v10 on data combined from multiple experiments as detailed in figure legends. Significance was defined as adjusted *p*<0.05.

### Data Availability

Raw .fastq and partially-processed RNAseq data will be deposited in the Gene Expression Omnibus at the time of publication. Values for all data points in graphs will be available at time of publication and reported in the Supporting Data Values file.

## Results

### Mice with transgenic IL-18 production show homeostatic NK- and CD8 T-cell changes reminiscent of Still’s disease

Given the association of high serum IL-18 and NK cytopenia/dysfunction in Still’s Disease and MAS^12,15,40,41^, we profiled NK cells and ILC1 in mice with transgenic expression of mature IL-18^36^. We observed a significant diminution of NK cells, particularly in livers and spleens, of *Il18tg* mice (Figure 1A). ILC1 were not significantly different (Supplemental Figure 1A). NK cells from *Il18tg* mice also showed a broad spectrum of maturity (as assessed by CD11b expression) and significantly lower expression of the IL-18 receptor (Figure 1B), consistent with our prior findings^33^ and Still’s patients’ NK cell IL-18 insensitivity^12,15^.

**Figure 1.**
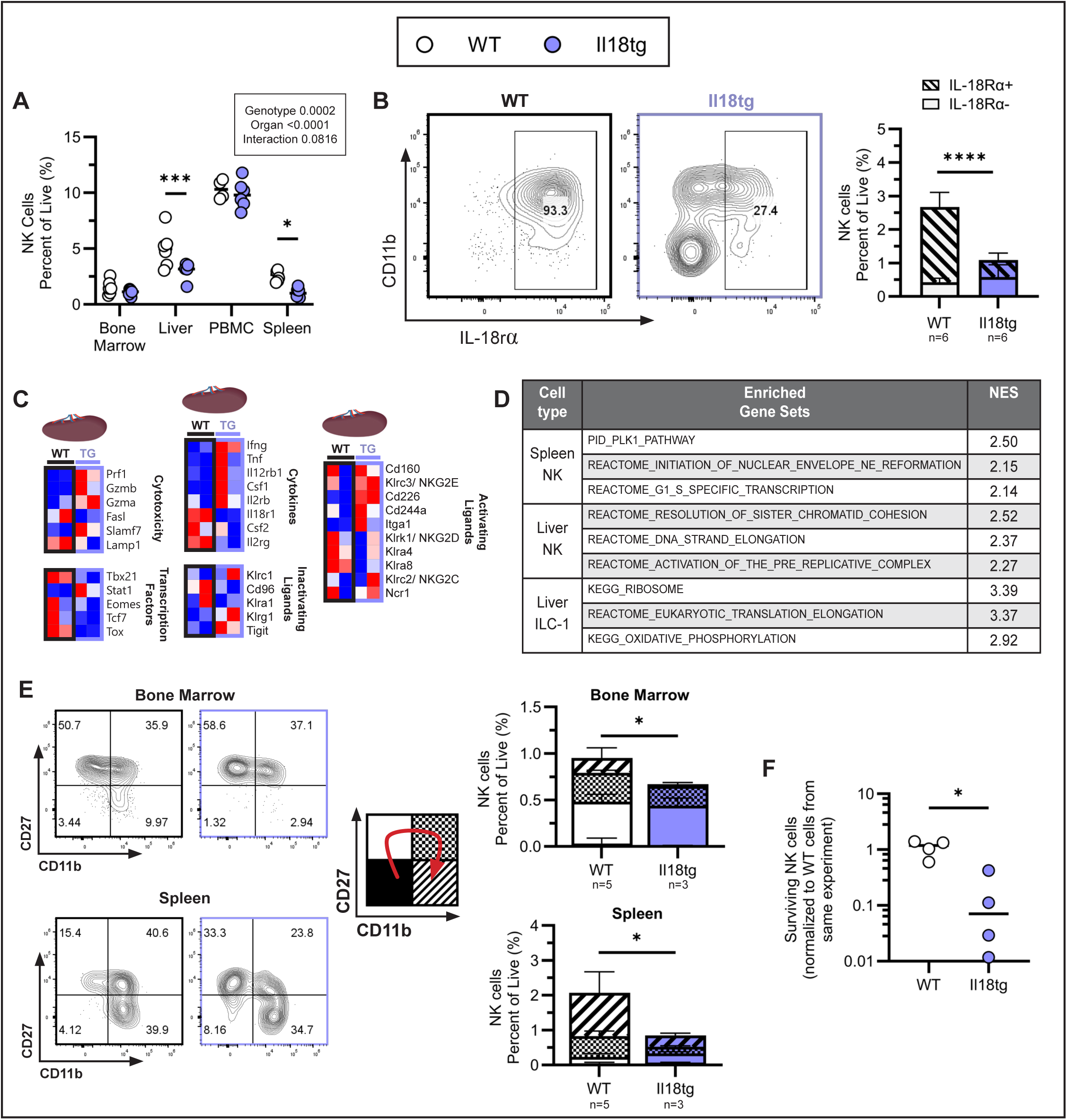
**Resting *Il18tg* mice harbor fewer and differently-activated NK cells** (A) NK cell proportion from various organs (NK1.1^+^ NKp46^+^ TCRb^-^ and CD127^-^ in spleen, TRAIL^-^ in liver) and (B) representative and aggregate splenic NK cell expression of CD11b and IL-18Rα. (C) Row-normalized splenic NK cell TPM values from selected canonical NK cell transcripts (see methods). (D) Top non-redundant gene sets enriched in *Il18tg* liver & spleen NK cells and liver ILC1 relative to WT (See also Supplemental Table 1). (E) Representative and aggregate data for NK maturity markers CD27 and CD11b on NK cells from bone marrow and spleen. Red arrow on represents the progression of NK cell maturation states. (F) Ratio of live NK cells after 5 hours in culture relative to live WT NK cells (average of WT technical replicates) from the same experiment. Each dot represents a biological replicate, combined data from two independent experiments. <u>Significance</u>: (A) 2-way ANOVA (box), with Šídák’s multiple comparisons test of genotype comparison within each organ and (B) unpaired t-test of IL-18Rα^+^ splenic NK cells. Representative of 4 independent experiments. (E) Unpaired t-test of mature (CD27^-^CD11b^+^) NK cells between genotypes. (F) Mann Whitney U-test. Only significant differences are shown. *=p<0.05, ***=p<0.001, ****=p<0.0001.

To better understand differences in these populations, we sorted spleen and liver NK cells and liver ILC1 directly ex vivo and performed bulk RNAseq. Differentially expressed gene (DEG) analysis highlighted downregulation of IL-18 receptor genes and the activating NK receptor NKG2D (*Klrk1*), and upregulation of GM-CSF (CSF2), but did not provide a clear explanation for their relative paucity or surface marker changes (Supplemental Figure 1).

Though some genes indicative of “type 1” immune activation were more abundant in NK cells from *Il18tg* mice (e.g *Prf1, Ifng, Klrc3, Cd226*), neither differentially expressed gene (DEG) analysis nor evaluation of the “Immunologic Signature” (MSigDB C7) gene sets showed consistent changes in canonical NK or ILC1-specific functions (Figure 1C, Supplemental Figure 1, Supplemental File 1). To probe more specifically for transcriptional changes associated with IL-18 exposure, we generated gene sets using RNA-seq data from primary murine splenic NK cells cultured with and without IL-18, IL-2, and/or IL-12 (Supplemental Methods, Supplemental File 2). We failed to detect significant enrichment of genes up- or down-regulated by ex vivo IL-18 in NK cells from *Il18tg* mice.

However, some “canonical pathway” (MSigDB C2) gene sets showed enrichment, particularly those related to transcription and replication (Figure 1D, Supplemental File 1). Finding fewer total NK cells in spleen and liver but not bone marrow, we hypothesized that NK cells in *Il18tg* mice were more immature. In both organs, we observed fewer cells with the CD27^-^CD11b^+^ phenotype of mature NK’s^42^ (Figure 1E). However, we did not detect enrichment of gene sets, generated from published RNAseq data^43^, associated with immature splenic NK, liver NK, or liver ILC1 cell transcriptomes (Supplemental Methods, Supplemental File 3).

Consistent with reports that IL-18 promotes human NK cell death in vitro^44^, we were prevented from assessing *Il18tg* NK cell function by their dismal survival ex vivo (Figure 1F). Contrasting with their NK cells, *Il18tg* mice showed an increase in spontaneous T-cell activation (CD44^+^PD1^+^) across all organs surveyed. This substantial increase in highly activated T-cells was almost entirely concentrated in the CD8^+^ compartment (Figure 2A-B, Supplemental Figure 2A-B). Thus, NK cells from unstimulated *Il18tg* mice are diminished in spleen and liver, have features of both transcriptional/replicative activity and immaturity, and poor ex vivo survival. Consistent with our prior findings^34^, their CD8 T-cells appear expanded and hyperactivated.

**Figure 2.**
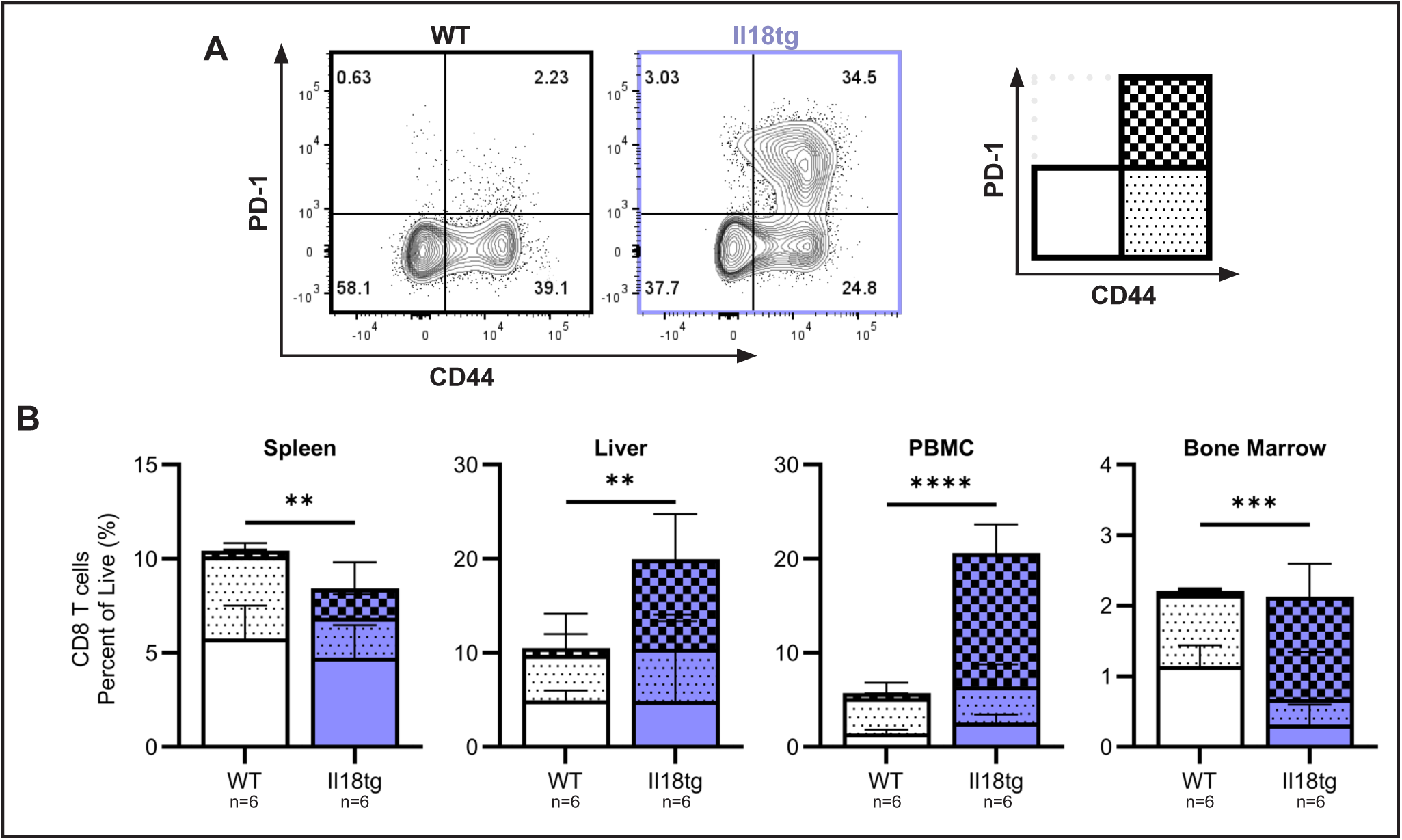
**Resting *Il18tg* mice demonstrate global increases in CD8 T-cell activation.** (A) Representative contour plots and (B) aggregate data for CD44 and PD-1 expression on splenic CD8 T-cells (Live, TCRb^+^CD4^-^CD8^+^) in resting mice. <u>Significance</u>: Unpaired t-test of highly activated (CD44^+^PD-1^+^) CD8 T-cells. Representative of 3 independent experiments. **=p<0.01, ***=p<0.001, ****=p<0.0001.

### Infection with mousepox virus (ectromelia) exacerbates spontaneous NK cell deficiency and CD8 T-cell hyperactivation

To assess NK function in vivo, we challenged *Il18tg* mice with the lytic DNA mousepox virus Ectromelia (ECTV, Moscow strain). Previously, we found that infection of *Il18tg* mice with the non-lytic, RNA-based Lymphocytic Choriomeningitis Virus (LCMV, Armstrong strain) induced cytokine storm, without compromising viral clearance, by hyperactivating CD8 T-cells^33–35^. However, NK cells contribute very little to the handling or clearance of LCMV^31,45^. Like LCMV, mousepox is typically cleared by WT C57BL/6 mice, but ECTV clearance relies much more heavily on the responsiveness and function of NK-cells, particularly at early timepoints^37^.

As expected, WT NK cells expanded and acquired activation/maturation markers (e.g. CD49b) 5 days after ECTV infection. However, NK cells from *Il18tg* mice showed no significant response to infection either numerically or by their activation state (Figure 3A-B). Despite this lackluster NK cell response, control of ECTV at the early timepoints associated with NK cell function was comparable between genotypes (Figure 3C). Despite defective NK responses, serum IFNg levels were comparable between WT and *Il18tg* mice (Figure 3D). CD8 T-cells, however, became even more activated at early timepoints after ECTV infection (Figure 3E). Given the contrast between diminished NK cell response to ECTV, but normal cytokine production and viral load, we predicted that NK/ILC1 depletion would have minimal effect on ECTV control in *Il18tg* mice relative to WT. Depleting mice of NK1.1-expressing cells prior to and during infection, we observed the expected rise in ECTV RNA in NK-depleted WT mice, but a much smaller effect in *Il18tg* mice (Figure 3F). Depletion showed a small increase in (already robust) CD8 T-cell activation in *Il18tg* mice, and did not affect systemic IFNg levels in either genotype (Figure 3G-H).

**Figure 3.**
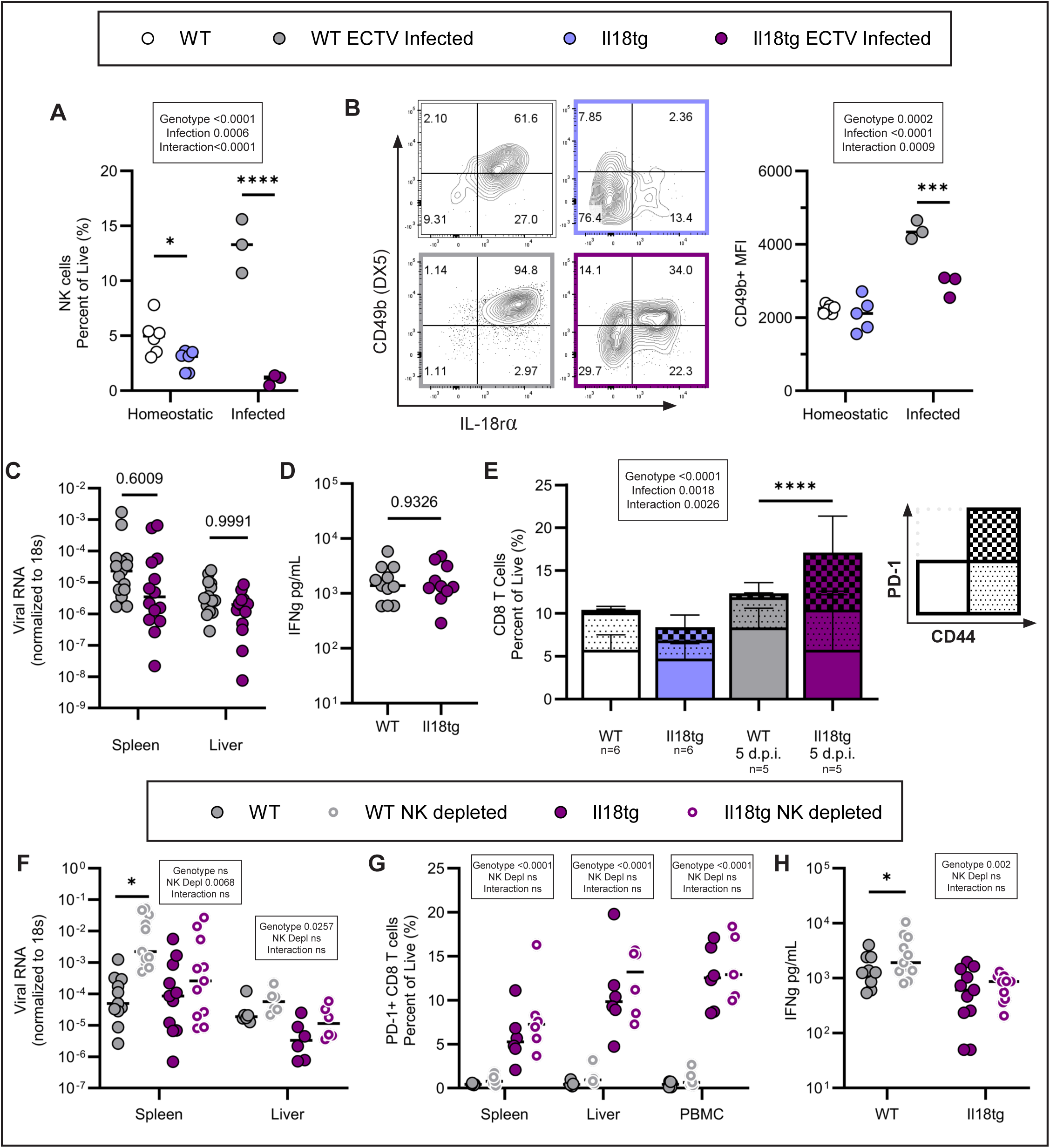
**Defective NK cell responses, but CD8 T-cell hyperactivation and normal viral control in *Il18tg* mice early after mousepox (ECTV) infection** Adult mice were infected with mousepox (ectromelia, ECTV) and assessed on Day 5 post-infection for NK cell proportion (A); CD49b and IL-18Rα expression (B); Viral RNA (C, see also **Supplemental** Figure 3); blood IFNg (D); and splenic CD8 T-cell activation (E). Mice were infected similarly and treated NK cell depleting or control antibodies (Days −1 & +2) and assessed at Day 5 for viral RNA (F); CD8 T-cell activation (G), and blood IFNg (H). <u>Significance</u>: 2-way ANOVA with Šídák’s multiple comparisons test of genotype (A-C, E) or NK depletion (F-H); or unpaired t-test (D). (A&B) Data representative of 2 independent experiments, and (C-H) combined from 2 independent experiments. *=p<0.05, ***=p<0.001, ****=p<0.0001.

### Il18tg mice ultimately develop fulminant systemic hyperinflammation following ECTV infection

As expected, WT mice demonstrated modest weight loss beginning around Day 6-7 post infection. Though *Il18tg* mice appeared similarly well to WT mice prior to Day 6, they quickly deteriorated with weight loss, anemia, thrombocytopenia, hepatic and splenic infarcts, profound hepatitis, and ultimately a high degree of morbidity (Figures 4A-E). In WT mice, liver and spleen inflammatory cells followed a predictable pattern, with early NK expansion yielding to CD8 T-cell activation that peaked around day 7-8. Myeloid cells, predominantly monocytes/macrophages, expanded modestly over time (Figures 4F-H). *Il18tg* mice, by contrast, failed to expand their NK pool but instead showed early and profound CD8 T-cell activation that was sustained over time. Though *Il18tg* mice handle LCMV without problems^33–35^, their handling of ECTV diverged in spleens by Day 6 and in liver soon thereafter (Figure 4I). As with CD8 T-cell hyperactivation, serum IFNg rose earlier and stayed elevated longer in *Il18tg* mice (Figure 4J). Thus, at later timepoints, *Il18tg* mice succumbed to disease with features of both immunodeficiency and hyperinflammation.

**Figure 4.**
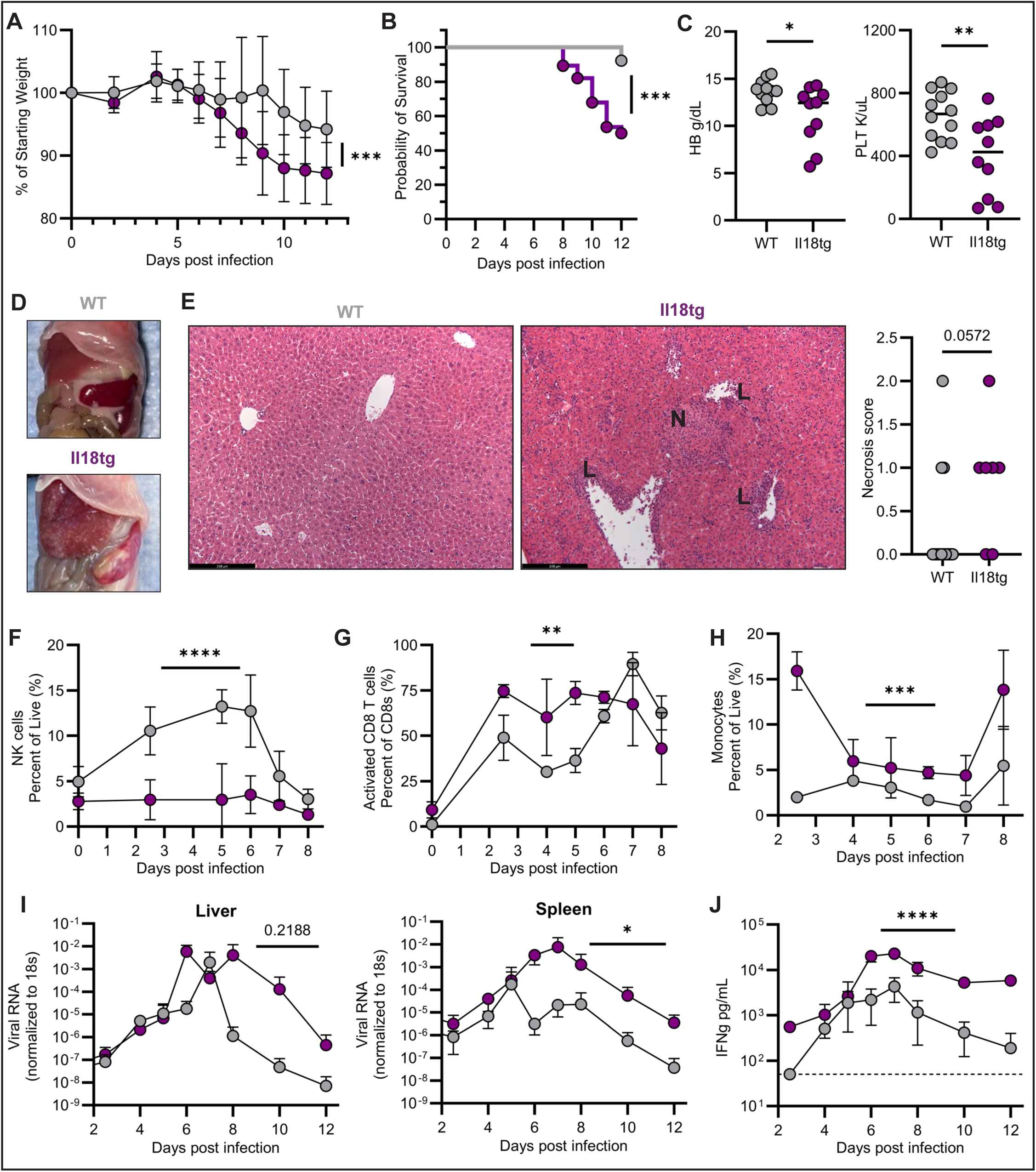
**Mousepox ultimately induces fatal HLH/MAS-like immunopathology and immunodeficiency in *Il18tg* mice** Adult mice were infected with mousepox and assessed for daily weight loss (A) and survival (B), (n=26 WT and 28 *Il18tg*); Hemoglobin (HB) and Platelet count (PLT) (C), gross abdominal morphology (D), and liver necrosis score (E). L=lymphocytic infiltration and N=area of necrosis. Mice were also assessed for liver NK (F) and activated CD8 T-cells (G), splenic monocytes (H) (CD11b^high^ SSC-A^low^ TCRb-B220-NK1.1-), liver and spleen ECTV RNA (I), and serum IFNg (J) at daily intervals post-infection; n= 2-5 per day. Dotted line in (J) represents threshold of detection. <u>Significance</u>: (A, F-J) Comparison of outcome variable over time between genotypes using Mixed Effects Model, (B) Kaplan-Meier test, (C) unpaired t-tests, and (E) Mann Whitney U-test (E). (A-E, I-J) Data combined from 4 independent experiments, (F-H) representative of 4 independent experiments. *=p<0.05, **=p<0.01, ****=p<0.0001.

### Il18bp^KO^ mice also develop severe HLH/MAS-like immunopathology and immunodeficiency with mousepox infection, despite normal basal NK state

We hypothesized that the apparent NK immunodeficiency in ECTV-infected *Il18tg* mice was due to acute versus chronic exposure to excess IL-18. IL-18 Binding Protein (IL-18BP) is an abundant, soluble, IFNg-induced antagonist that binds to activated IL-18 with high affinity and prevents it from signaling^21,46^. In our experience, unstimulated *Il18bp*^KO^ mice show no elevation in systemic IFNg^47^, and we observed little effect on NK cell number or CD11b, CD49b, or IL-18R expression (Figure 5A, Supplemental Figure 1B). Despite their more normal start, NK cells from *Il18bp*^KO^ mice also failed to expand or upregulate CD49b appropriately upon mousepox infection (Figure 5B). In contrast to *Il18tg* mice, *Il18bp*^KO^ mice already showed a significant elevation of ECTV RNA at Day 5 post-infection (Figure 5C).

**Figure 5.**
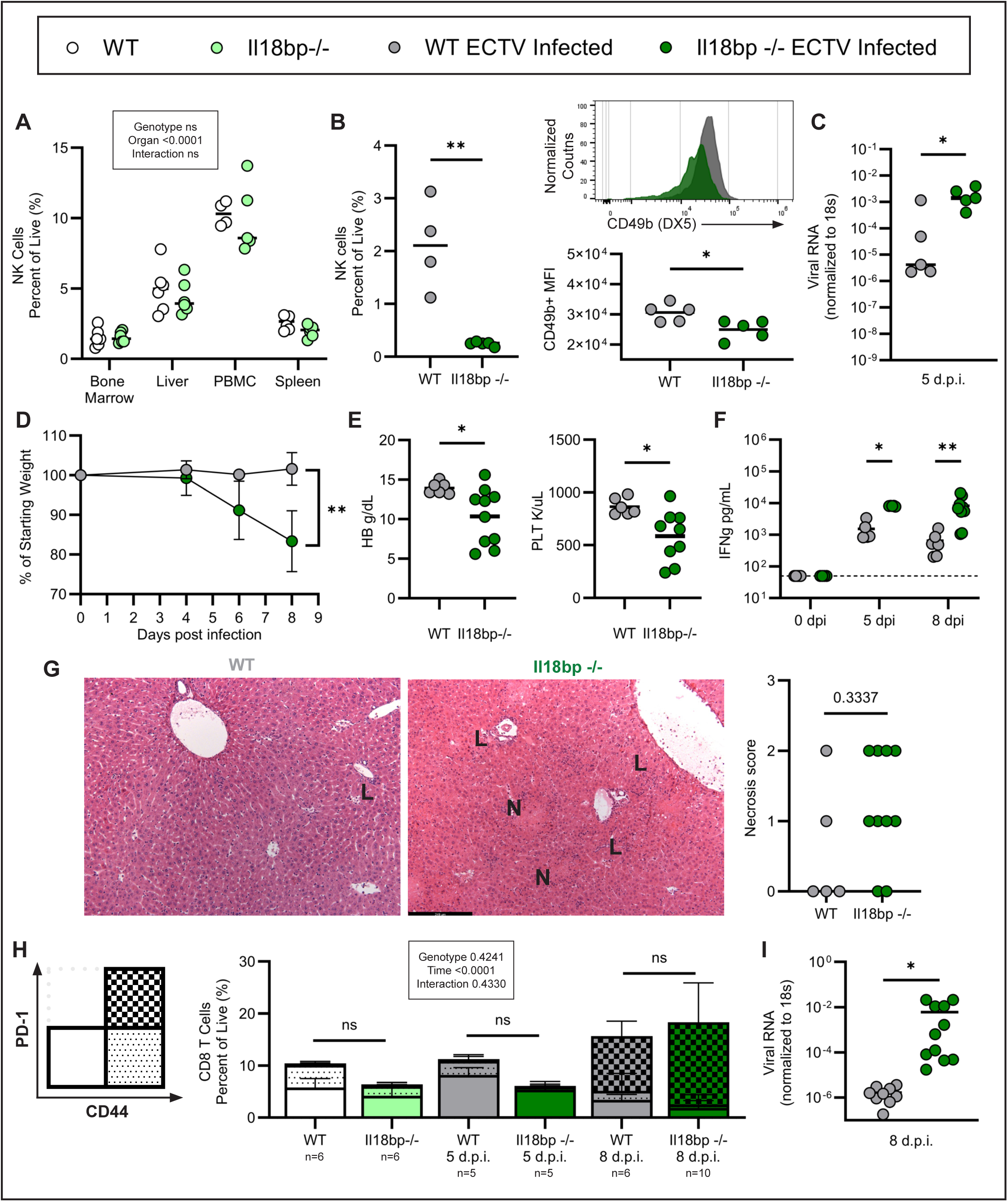
***Il18bp^KO^* mice develop severe HLH/MAS-like immunopathology and immunodeficiency with mousepox infection, despite normal basal NK state.** Homeostatic NK cell proportions from various organs (A). Adult mice were infected and assessed 5 days post mousepox infection for splenic NK cell frequency and CD49b expression (B) and splenic viral load (C). Daily weight (D); day 8 hemoglobin (HB) and platelet count (PLT) (E), serum IFNg (F), liver hepatitis/necrosis (G), frequency of activated splenic CD8 T-cells (H), and day 8 splenic ECTV RNA (I). Dotted line in (F) represents threshold of detection. L= lymphocytic infiltration and N=necrosis (G). <u>Significance</u>: 2-way ANOVA with Šídák’s multiple comparisons posttest comparing genotypes within each organ (A) or timepoint (F,H), unpaired t-test (B, C, E, I), mixed-effects model comparing genotypes over time (D), or Mann-Whitney test (G). Only CD44^+^PD1^+^ CD8 T-cells were compared in (H). Data representative of 2 independent experiments. *=p<0.05, **=p<0.01.

At later timepoints, *Il18bp*^KO^ mice exhibited similar weight loss, cytopenias, IFNg elevation, hepatitis, and prolonged viremia as *Il18tg* mice (Figure 5D-G). Notably, *Il18bp*^KO^ mice showed no significant differences in CD8 or CD4 T-cell activation at any timepoint, although CD44^+^PD-1^+^ CD8 T-cells trended toward greater expansion in infected *Il18bp^KO^* mice by later timepoints (Figure 5H, Supplemental Figure 2). Despite a less profound degree of CD8 T-cell activation, *Il18bp*^KO^ mice showed much higher IFNg elevation and collapse of hepatic architecture, with wide swaths of necrosis and early fibrosis (Figure 5F, Supplemental Figure 4).

As it is possible that IL-18BP has as-yet undescribed functions beyond restraining IL-18, we challenged mice deficient in both IL-18 and IL-18BP. Fatal ECTV in *Il18bp*^KO^ mice appeared to follow directly from unopposed IL-18, as *Il18^KO^Il18bp*^KO^ mice resisted HLH and handled ECTV similarly to WT mice (Supplemental Figure 5). Though *Il18tg* mice appear transiently protected (relative to *Il18bp*^KO^ mice) at very early timepoints, these data suggest that excess IL-18, whether due to chronic IL-18 overproduction (*Il18tg*) or failure to oppose acutely induced IL-18 (*Il18bp*^KO^), leads to lethal responses following mousepox infection.

### Excess IL-18 induces an ineffective CD8 T-cell response following mousepox infection

Even without appropriate NK responses, we wondered why rapid and robust CD8 T-cell activation in ECTV-infected *Il18tg* mice was not sufficient to curtail fatality. To better understand the nature of their pathogen-specific responses, we tracked ECTV-specific CD8 T-cells using tetramer staining for the immunodominant B8R^20–27^ ECTV peptide^48^. Whereas WT mice developed the anticipated, robust expansion of B8R-specific CD8 T-cells, expansion of this population in *Il18tg* livers and spleens was both delayed and far less pronounced (Figure 6A).

**Figure 6.**
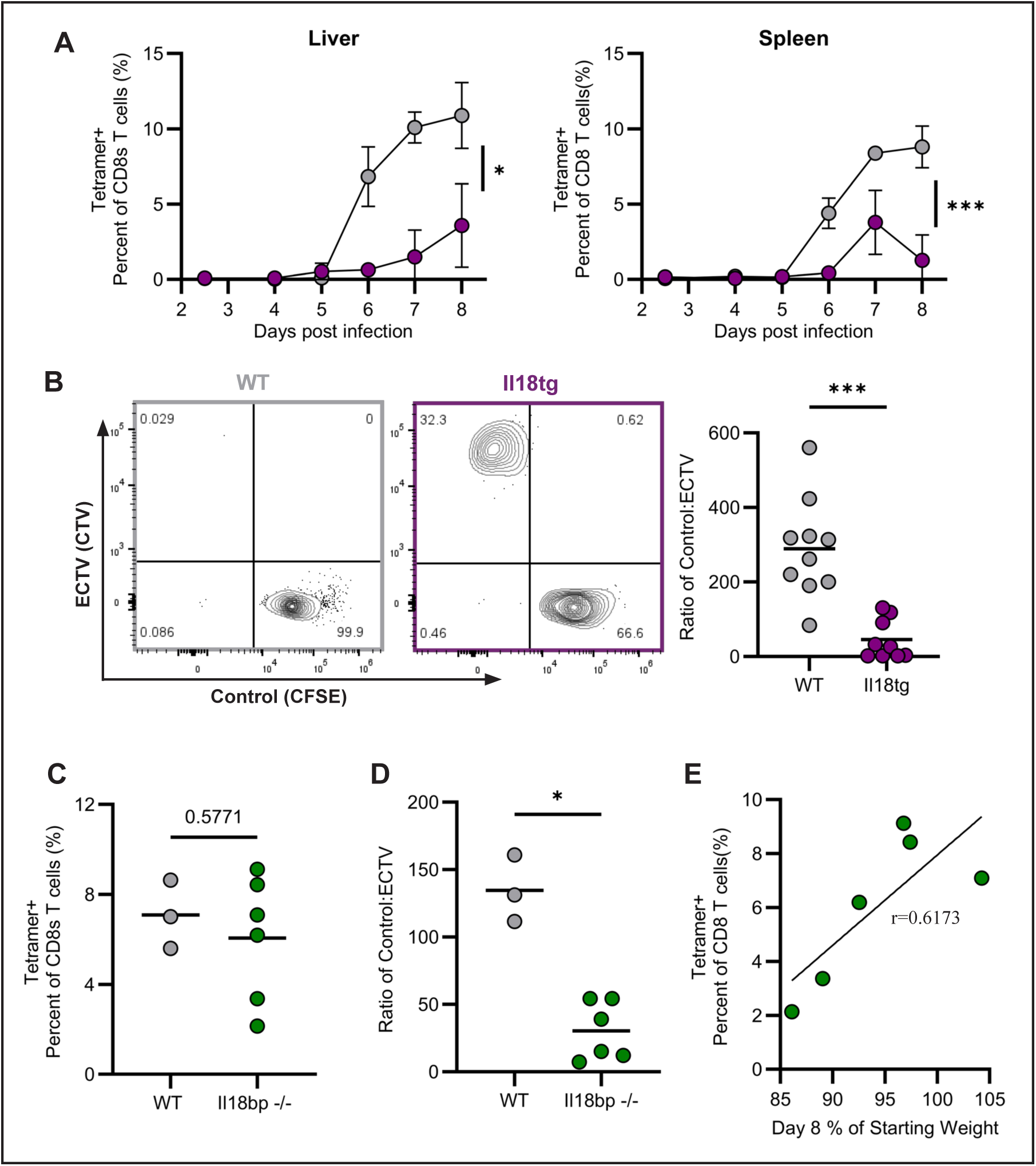
**Failed antigen-specific CD8 T-cell expansion and function in mice with excess IL-18 following mousepox infection.** Mice of the indicated genotypes were infected with ECTV and evaluated periodically (A) or only on day 8 (C) for CD8 T-cells specific for the immuno-dominant ECTV BR8 peptide. In vivo killing assay in ECTV-infected mice of the indicated genotypes (B, D): the ratio of control splenocytes (CFSE-labeled) to BR8 peptide-loaded splenocytes (CTV-labeled) 4 hours after transferring at 1:1 ratio. (E) Correlation between weight loss and BR8-tetramer^+^ CD8 T-cells recovered in spleens on day 8 amongst *Il18bp^KO^* mice. <u>Significance</u>: mixed effects model comparing genotypes over time (A), unpaired t-test (B-D), and linear regression (E). Data representative of 3(A) and 2(C-D) independent experiments and (B) combined from 2 independent experiments. *=p<0.05, ***=p<0.001.

Similarly, an in vivo killing assay^49^ showed that infected WT mice specifically eliminated splenocyte targets coated with the ECTV-peptide, whereas infected *Il18tg* mice showed dramatically impaired clearance of BR8-coated targets (Figure 6B). ECTV-tetramer+ CD8 T-cells were more variably abundant in spleens of *Il18bp*^KO^ mice, but they nevertheless failed to clear peptide-coated cells (Figures 6C-D). In individual *Il18bp*^KO^ mice, disease severity correlated with B8R-specific CD8 T-cells (Figure 6E).

### Mousepox-specific CD8 T-cells fail to expand in Il18tg mice despite early and sustained CD8 T-cell activation

To better understand why *Il18tg* mice failed to mount an effective CD8 T-cell response, we examined the longitudinal characteristics of CD8 T-cells pulled from 10,000 live cells per organ (liver and spleen), per mouse, per day by flow cytometry (Figure 7A). There was substantial overlap between liver and spleen CD8 T-cells across genotypes and activation markers (Figure 7B,C). WT mice develop a population of highly activated CD8 T-cells around 7-8 days post-infection, including those that bound the B8R-tetramer, marking them as pathogen-specific effector CD8 T-cells. *Il18tg* mice showed significant populations of hyperactivated CD8 T-cells as early as 2.5 days post-infection, suggesting activation of cells not responsive to ECTV-antigens. The characteristics of their CD8 T-cell pool shifted inconsistently over time, but very few *Il18tg* CD8 T-cells cells acquired tetramer staining at any timepoint. A small population of PD-1^+^CD8 T-cells with low CD44 expression and no B8R-tetramer binding was present in *Il18tg* mice, reminiscent of a similar population observed as *Il18tg;Prf1^KO^* mice develop spontaneous HLH/MAS^34^ (Figure 7C,D). The few B8R+ CD8 T-cells that arose in *Il18tg* mice did not appear until day 8, but otherwise bore normal activation marker expression. These data suggest impaired antigen-specific CD8 T-cell development/expansion in *Il18tg* mice, despite early and continuous CD8 T-cell hyperactivation.

**Figure 7.**
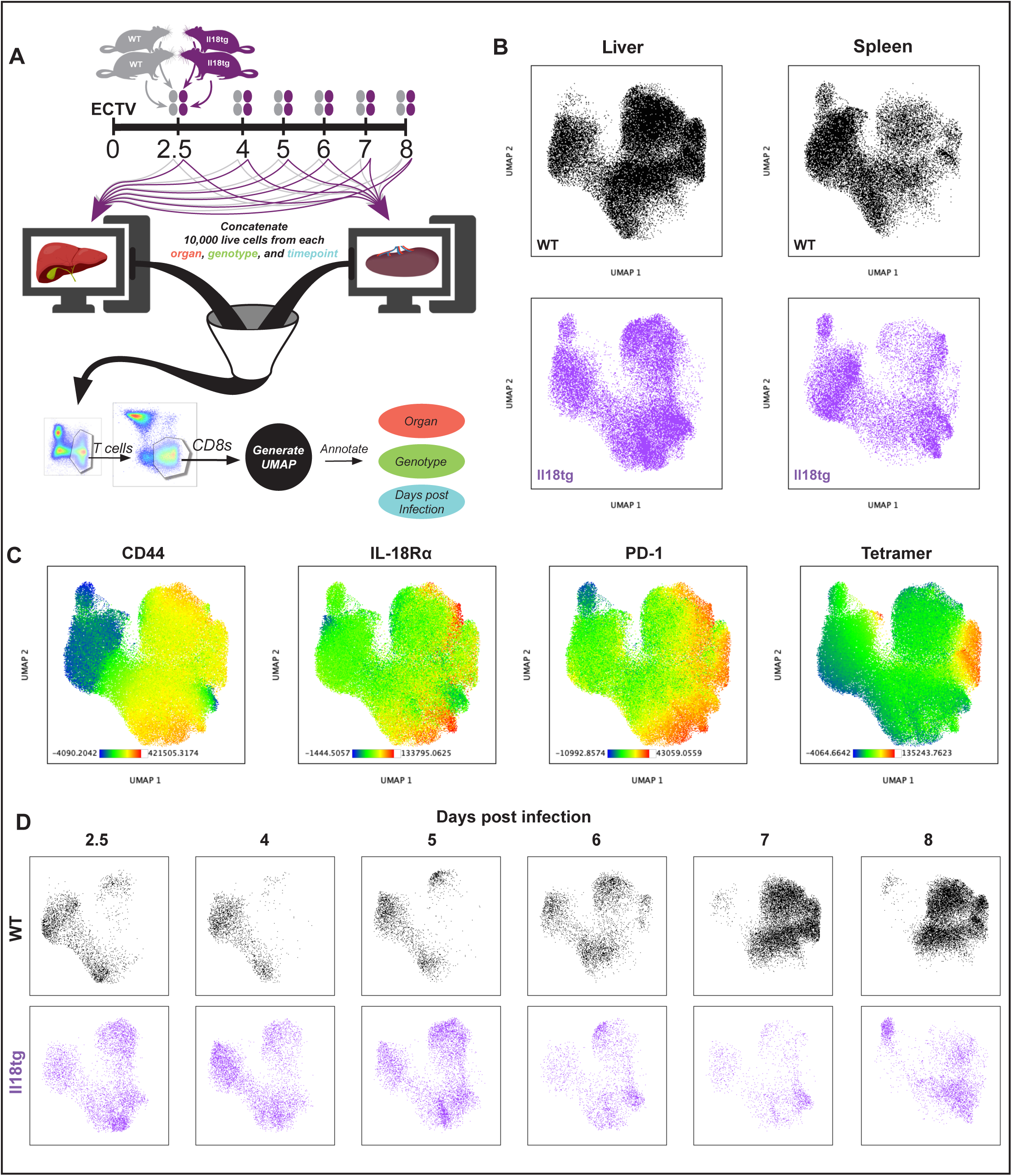
**Early and persistent CTL hyperactivation belie failed antigen-specific CD8 T-cell responses in mousepox-infected *Il18tg* mice.** WT and *Il18tg* mice were infected with ECTV and livers and spleens from two mice/genotype were evaluated by flow cytometry at the indicated timepoints post-infection. 10,000 live cells per organ from each mouse were concatenated, and from this CD8 T-cells were gated and used to generate the annotated UMAP (A). UMAP distribution of cells by organ and genotype (B). Fluorescence intensity of the indicated surface molecules (C). Progression of CD8 T-cell abundance and location within the UMAP over time and by genotype (D). Parameters used to generate UMAP are described in Methods.

#### Activated NK cells protect Il18tg mice from ECTV-induced immunopathology

Effective NK and antigen-specific CD8 T-cell responses are required sequentially for protection from mousepox^50,51^. Both *Il18tg* and *Il18bp^KO^* mice showed NK and CD8 T-cell defects following mousepox infection, but not LCMV infection^33^. We hypothesized that WT NK cells would be sufficient to promote effective pathogen-specific responses and protect *Il18tg* mice from disease. Indeed, transfer of WT NK cells expanded ex vivo enabled *Il18tg* mouse survival, viral clearance, and antigen-specific CD8 T-cell responses similar to WT (Figure 8A-C). IL-18 receptor expression was intact (Figure 8D), suggesting that the acute effects of excess IL-18 on pre-activated NK cells are not sufficient to impair their function and allow fulminant HLH/MAS.

**Figure 8.**
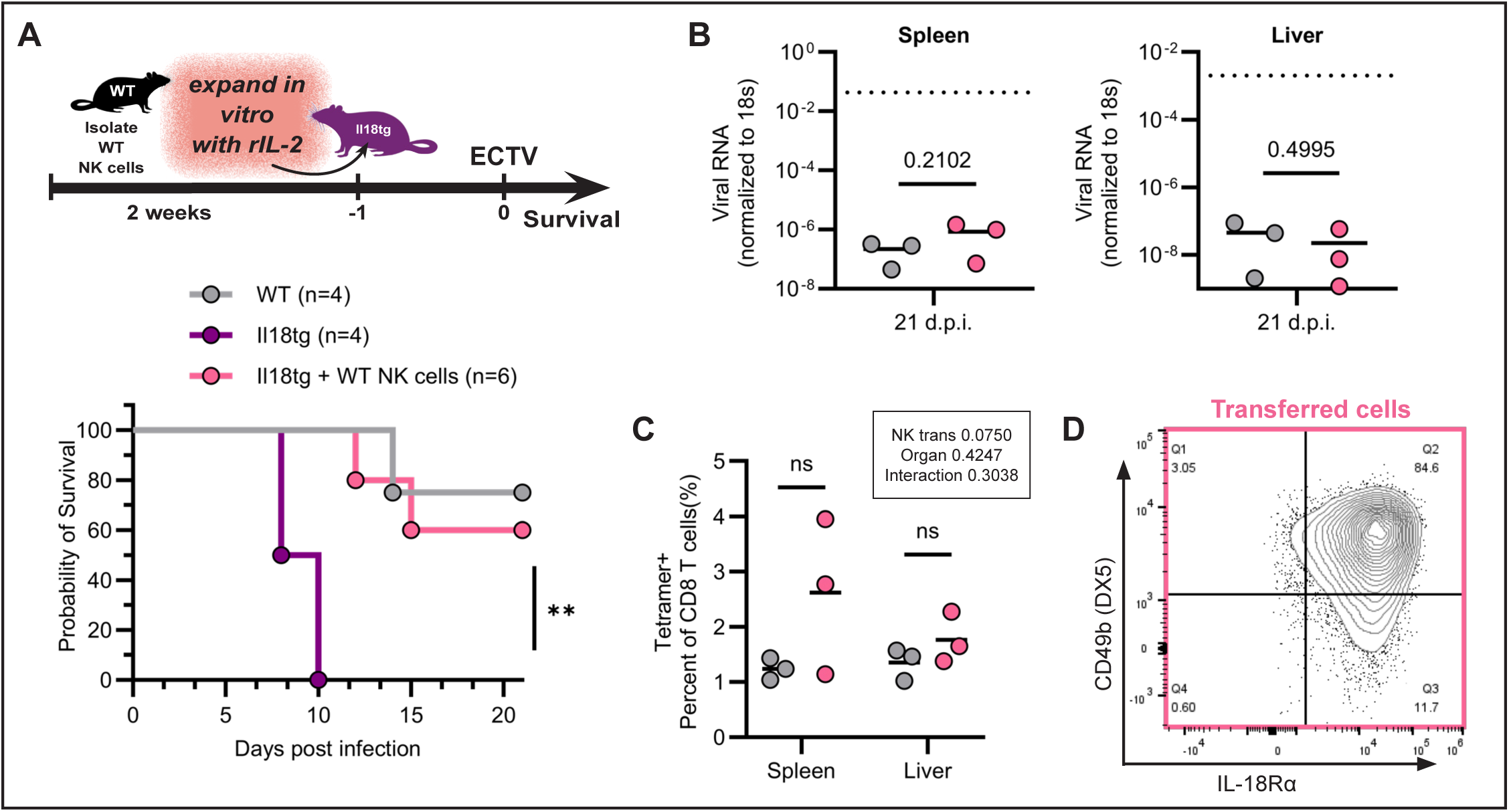
**Adoptive transfer of NK cells preactivated in vitro rescues *Il18tg* mice from ECTV pathology.** (A) Experimental design and survival of *Il18tg* mice following adoptive transfer of pre-expanded NK cells one day prior to infection. (B) ECTV RNA from organs collected from mice 21 days post infection. Dotted line represents mean value of *Il18tg* mice 8-12 days post-infection. (C) ECTV BR8-tetramer^+^ CD8 T-cells 21 days post infection. (D) Representative contour plot of CD49b and IL-18Rα expression of pre-expanded NKs just prior to adoptive transfer. <u>Significance</u> was established using (A) Kaplan-Meier test of *Il18tg* vs *Il18tg*+NK cells, (B) unpaired t-test, (C) 2-way ANOVA with Šídák’s multiple comparison test of WT versus Il18tg + WT NK cells within organs. **=p<0.01.

## Discussion

HLH is a complex life-threatening systemic inflammatory state, conceptually and mechanistically overlapping with sepsis and Cytokine Release Syndrome. Though identifying patients “in HLH” is important for risk assessment, monitoring, and establishing a diagnostic heuristic, the most important task for clinicians managing HLH is identifying/targeting HLH contributors. This task is not simple. Not all contributors are known, and some are difficult to identify (e.g. occult lymphoma). Additionally, they do not act in isolation. FHL’s enigmatic variation in age at presentation is likely related to its basis in the interaction of defective GMC and an infectious trigger.

Finally, individual contributors may have multiple mechanisms of immunopathology: *autoinflammation* involves excessive innate immune responses; *immunodeficiency* involves direct pathogen-induced damage due to impaired control; and we define *hyperinflammation* as damage from excessive, pathogen-specific, adaptive immune responses. We previously found that IL-18, a uniquely specific biomarker of MAS susceptibility^18,19^, drives HLH/MAS by *autoinflammatory* IFNg amplification^18,47^, synergy with GMC impairment^33,34^, and promotion of inflammatory target cell death^35^; and by *hyperinflammatory* CD8 T-cell activation^33^. Herein, we describe a new, context-specific role for IL-18 as an inducer of NK *immunodeficiency*.

Even without MAS, Still’s patients often have free IL-18 elevation^19^, their normal peripheral NK cell number belie their abnormal NK responses. Transcriptionally, their NK cells have upregulated innate immune genes like *TLR4* and *S100A9*, and functionally they show specific defects with IL-18 responsiveness^12,15,16^ (recapitulated in *Il18tg* mice^33^, Figure 1B).

Active MAS patients show peripheral NK cytopenia but their tissues show lymphohistiocytosis^52^. NK cytopenia is common in most forms of secondary HLH^53^ or even in severe COVID-19^54^, and may correlate with a shared HLH/MAS immunophenotype of CD8^+^ T-cell and macrophage hyperactivation^52,55–58^. Experimental evidence supports this shift from NK to CD8 T-cell activation: both IL-18 and IL-6 appear to impair NK cell proliferation and survival^14,44^, and activated CD8 T-cells serve as a sink for a key NK proliferation/survival signal IL-2^59^.

Profound IL-18 elevation may be under-recognized in IA-HLH. HLH/MAS arises in most patients with XIAP-deficiency, typically triggered by EBV infection. Though XIAP deficiency may promote inflammasome activation in experimental systems, the mechanistic basis for these patients’ highly-elevated IL-18 and susceptibility to EBV is unknown^24,60,61^. Excess IL-18 may also promote viremia and hyperinflammation in Kaposi’s sarcoma herpesvirus (KSHV)-associated inflammatory disorders^62,63^. Two recent EBV-HLH patients further illustrate this point. Patient 1 showed the marginal IL-18 elevation expected in EBV-HLH, despite a severe course. Patient 2 was found to have profound IL-18 elevation at presentation that persisted through discharge (Table 1). Both were previously healthy, lacked clinical features of Still’s Disease, showed no genetic susceptibility to EBV or HLH on exome sequencing, and ultimately responded well to immunomodulation. We also found MAS-level IL-18 elevation in a small proportion of patients from infection-associated HLH^18^ and hyperferritinemic sepsis^64^ cohorts. Highly-elevated IL-18 levels may be under-detected in other infection cohorts due to widespread hook effects in multiplex panels^18,64^.

**Table 1.**

| Table 1 | NI range | Patient 1 (3y) |  |  | Patient 2 (14y) |  |  |
| --- | --- | --- | --- | --- | --- | --- | --- |
|  |  | 1st | Worst | DC, d18 | 1st | Worst | DC, d10 |
| ALT | < 26 U/L | 175 | 936 | 95 | 382 | 945 | 708 |
| LDH | 146-386 U/L | 3194 | " | ND | 5335 | " | 958 |
| CRP | <0.9 mg/dL | 6.3 | 8.6 | 0.5 | 2.9 | " | 0.5 |
| ferritin | 10-99 ng/mL | 39041 | 79373 | 1064 | 55505 | 59560 | 8127 |
| sIL-2Ra | <858 pg/mL | 63424 | " | 1279 | 17564 | " | 6264 |
| IFNγ | < 6.5 pg/mL | 11,500 | NA | NA | 4720 | " | 297 |
| CXCL9 | < 647 pg/mL | 36223 | " | 180 | 211105 | " | 70557 |
| IL-18 | < 477 pg/mL | 4243 | " | ND | 368966 | " | 186439 |
| EBV | 0 IU/mL | 5000000 | " | 367504 | 360181 | " | 4859 |
| ANC | 1600-8300 /uL | 2260 | 360 | 1160 | 6620 | 1450 | 2070 |
| **Hgb** | 11.5-16 g/dL | 7 | " | 9.4 | 11 | 8.2 | 9.4 |
| **Plt** | 150-450 EE3/uL | 18 | 15 | 758 | 118 | 87 | 333 |
| **fibrinogen** | 131-405 mg/dL | 130 | 54 | 152 | 70 | 67 | 156 |
| **Immunomodulation** |  | GC, IL-1RA, anti-IFNg,<br>etoposide |  |  | GC, IL-1RA |  |  |
| DC=at discharge; ALT=Alanine aminotransferase; LDH=lactate dehydrogenase; ND=not done; CRP=C-reactive protein; sIL-2Ra=soluble IL-2 receptor alpha; IFNg=Interferon gamma; NA=not applicable due to use of neutralizing antibody; ANC=absolute neutrophil count; Hgb=hemoglobin; Plt=platelet count; GC=high-dose glucocorticoids; IL-1RA=IL-1 Receptor Antagonist anakinra. Black background = lower in HLH. |  |  |  |  |  |  |  |

Our observations in *Il18tg* mice illustrate how NK-mediated protection extends beyond early IFNg production and GMC. Like MAS patients, *Il18tg* mice showed a basal state of mild NK cytopenia, CD8 T-cell activation, and elevated IFNg. Strangely, this did not correlate with a clear transcriptional program of immaturity, but rather activation/replication activity. NK cells in *Il18bp*^KO^ mice appeared more like WT under resting conditions, though more significant basal NK defects have been observed in *Il18bp*^KO^ mice in other facilities^65^. *Il18tg* NK cells struggled to survive in ex vivo cultures (Figure 1F) and their expression of integrins like αM and α2 (CD11b and CD49b, respectively), was more impaired than their expression of critical activating receptors like NKG2E and NKG2D. This suggests that IL-18 impaired NK cell viability and in vivo proliferation greater than per-cell function^51,66,67^, a pattern indicative of activation-induced cell death (AICD) rescuable by transfer of “pre-activated” NK cells. These baseline defects engendered by IL-18: poising NK cells for AICD and amplifying homeostatic CD8 T-cell activation, were fatally exploited by mousepox.

*Il18tg* mice contained ECTV normally until day 5, despite inert NK responses, likely due to IFNg produced from antigen non-specific CD8 T-cells. *Il18bp^KO^* mice, by contrast, quickly acquired the problems present constitutively in *Il18tg* mice. The benefit of a pre-existing effector CD8 T-cell pool was merely a weak and temporary restraining of ECTV for the first five days after infection. Both strains then failed to mount effective pathogen-specific CD8 T-cell responses. In this respect, IL-18 joins other NK defects (e.g. loss of NKG2D, virulence factor inhibition of NK immune synapse formation, and aging-related migration defects) in demonstrating that, specifically for lytic DNA viruses like mousepox and murine cytomegalovirus (MCMV), robust NK responses are a requirement for effective antigen-specific CD8 T-cell responses^37,51,68,69^.

This study has several important limitations. Mousepox is an excellent in vivo NK challenge, but not a human pathogen. Efforts to develop EBV-based models of HLH are ongoing^70–72^. Unlike XIAP deficiency, prolonged EBV viremia is not commonly reported in MAS, although viremia in primary infection can last months and local reactivation is common even in normal hosts^73^. IL-18 mediated NK immunodeficiency may be local/transient and serve to further boost IL-18’s hyperinflammatory effects. Additionally, how NK cells prime effective CD8 T-cell responses to lytic DNA viruses like ECTV and MCMV remains a mystery. The robust non-specific CD8 T-cell activation observed in *Il18tg* mice suggests NK cells are not required for activation in general. Likewise, the absence of a significant pool of pathogen-specific cells at any timepoint argues against exhaustion. Other hypotheses, like NK effects on antigen presentation, fractricide, and direct protection from organ dysfunction, should be explored.

Overall, this study shows how one cytokine may contribute to HLH via multiple simultaneous mechanisms depending on the presence/nature of infectious trigger. IL-18’s auto/hyperinflammatory propensity for CD8 T-cell activation likely combines with its promotion of NK activation-induced cell death (AICD) to drive NK immunodeficiency in MAS. Even using murine models, specific stimuli, longitudinal quantitation of viral load, and detection of tetramer-labeled T-cells, we struggle to apportion immunopathology amongst IL-18’s mechanisms. In HLH patients, attribution to one mechanism versus another might dramatically affect treatment strategy, but is often practically impossible. The recent regulatory approval of IFNg blockade for steroid-refractory MAS represents major progress^74^, but not all MAS patients respond favorably.

Neither acute IFNg nor IL-18 blockade will be effective if the wheels of NK immunodeficiency and/or CD8 T-cell hyperinflammation are already in motion. No IL-18 blocking drugs are yet approved for clinical use, although multiple strategies are in development and anecdotes have been encouraging^2^. Our findings support assessment of IL-18 in HLH patients as part of a broad assessment of likely contributors. However, they undermine the notion that HLH can be successfully treated by targeting immune responses alone, underscoring individualized treatment paradigms that balance immunomodulation with aggressive detection and clearance of contributing pathogens.

## ACKNOWLEDGMENTS

the authors would like to acknowledge Giuseppe Sciume (Sapienza University, Rome) for assistance with in vitro NK cell RNAseq analysis; Elise Peauroi for helpful advice; Rania Elbakri, William MacDonald, and Amanda Poholek (University of Pittsburgh) for assistance with RNAseq library prep; Florin Tuluc and Jennifer Murray (CHOP Flow Cytometry Core Laboratory, RRID:SCR_009726) for assistance with Flow Cytometry experiments, the CHOP Research Institute Pathology Core for assistance with histology imaging, and the NIH Tetramer Core Facility for key reagents.

## Conflict-of-interest statement

The authors have declared that no conflict of interest exists.

## Notes

### Competing Interest Statement

The authors have declared no competing interest.

