## Supplemental Figures for "Multimodal induction of fulminant HLH by IL-18 includes virus-specific NK immunodeficiency"

**Supplemental Figure Legends:**

**Supplemental Figure 1. Altered NK cell phenotype in resting *Il18tg* more than *Il18bp*$^{KO}$ mice.** Companion to Figure 1. (A) ILC1 cells from spleen (NK1.1+ NKp46+ CD127+) and liver (NK1.1+ Nkp46+ CD49b- TRAIL+). (B) Representative and aggregate homeostatic NK cell CD11b, CD49b, and IL-18Rα expression from NK cells. (C) Row- and organ-normalized TPM values from selected canonical NK cell transcripts (see methods) from Bulk RNAseq of sorted Liver NK cell and ILC1. Significance: 2-way ANOVA with Tukey's pairwise post-test between genotypes within each organ (A-B) Only significant differences of post-test are shown. Data representative of 3 independent experiments (A) and combined from 2 independent experiments (B). *=$p<0.05$, **=$p<0.01$, ***=$p<0.001$, ****=$p<0.0001$.

**Supplemental Figure 2. Excess IL-18 does not induce spontaneous CD4 T-cell hyperactivation.** Representative contour plots of splenic CD4 T-cell (Live, $TCRb^{+}CD4^{+}CD8^{-}$) expression of CD44 and PD-1 (A) and aggregate data from spleen, liver, PBMC, and bone marrow of the same mice (n=6). Significance: 2-way ANOVA with Tukey's pairwise post-test between genotypes within each cell type identified no significant post-test comparisons. Data representative of 3 independent experiments

**Supplemental Figure 3. Mousepox viral RNA correlates with PFU.** Viral RNA was measured via qPCR of Evm003 viral cDNA and normalized to 18s endogenous RNA. PFU determined via plaque assay of spleen from day 5, 8 and 12. $R^2$ value established by simple linear regression.

**Supplemental Figure 4. Trichrome staining revealed extensive necrosis in *Il18bp*$^{KO}$ livers.** Liver histopathology 8 days post infection visualized via trichrome staining. "N" denotes sites of necrosis.

**Supplemental Figure 5. Susceptibility of Il18bp$^{KO}$ mice to fulminant ECTV-HLH requires intact IL-18 production.** WT and *Il18*$^{KO}$*Il18bp*$^{KO}$ mice were infected with mousepox and assessed between days 8-12 for percent weight loss (A, n=4-5), hemoglobin and platelet count (B), CD8 T-cell activation and antigen-specific T-cell abundance (C&D), viral RNA from spleen (E), and plasma IFNg from 8-12 d.p.i (F). Significance: Mixed Effects Model comparing genotypes (A) and unpaired t-test (B-F). In (C), comparisons were restricted to PD-1+ CD44+ CD8 T-cells. Data representative of 2 independent experiments (A-B), combined from 2 independent experiments (D-F). **=p<0.01

**Supplemental Table 2. Chronic excess IL-18 induced cell cycling and replication transcriptional profile.** Bulk RNA sequencing data set was analyzed using GSEA (See Methods). Ranked gene lists were generated for each cell type for Il18tg vs WT comparison. GSEA results from comparison to canonical pathways curated geneset (C2) and immunologic signatures geneset (C7). Repeated core enrichment genes across genotypes are highlighted.

**Supplemental Table 3. Excess IL-18 ex- vivo alters NK cell transcriptional prolife, does not resemble chronic excess IL-18 exposure**. Custom genesets were produced using previously published and ammened bulk-RNA sequencing dataset of ex vivo cytokine stimulated NK cells DGE list was generated for the comparison IL-18 alone & IL-18 + IL-12 versus no stim & IL-12 alone & IL-2 + IL-12. All differentially expressed genes with a log fold change > 2 and FDR p-value < 0.001 were used to generate up and down genesets. Enrichment of these genesets were assessed in Il18tg NK cells compared to WT NK cells using GSEA. Repeated core enrichment genes across genotypes are highlighted.

**Supplemental Table 4. Il18tg NK cells are not significantly enriched for immature genes.** Custom genesets were produced using previously published bulk-RNA sequencing dataset of progenitor, immature and mature NK cells (GSE77695). DGE lists were created comparing immature bone marrow NK cells versus mature splenic or liver NK cells and NK progenitors from the bone marrow versus liver ILC1 cells. All significant differentially expressed genes were used to generate genesets for each comparison. Enrichment of these genesets were assessed in Il18tg NK cells compared to WT NK cells using GSEA. Repeated core enrichment genes across genotypes are highlighted.

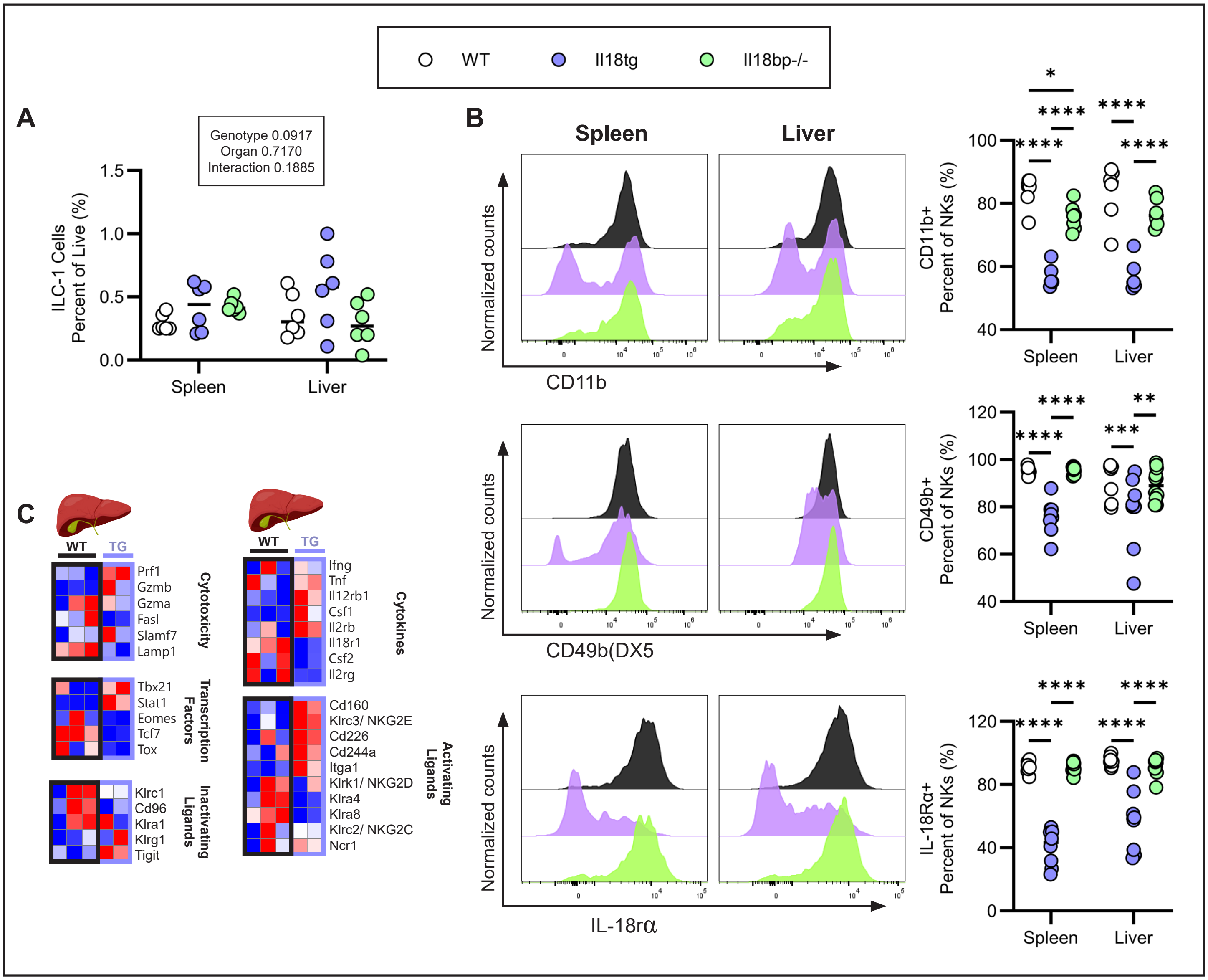


**Supplemental Figure 1**

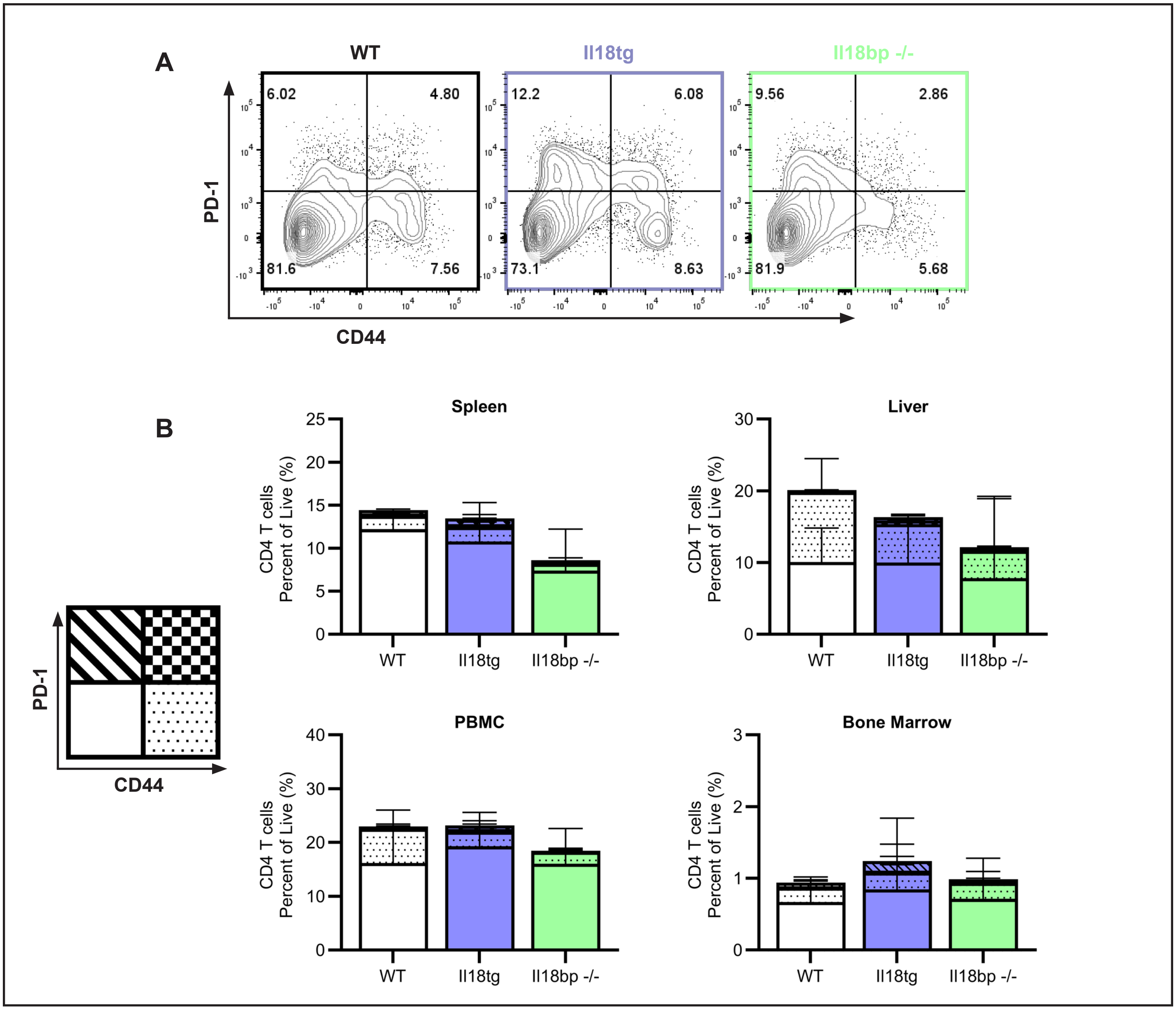


**Supplemental Figure 2**

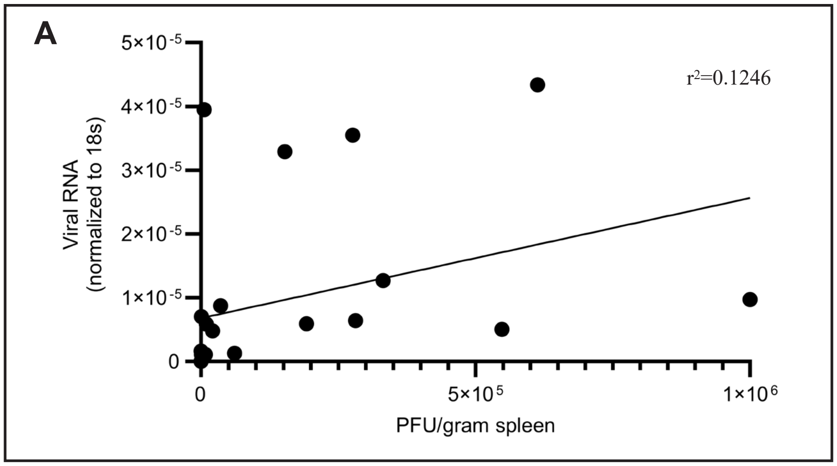


**Supplemental Figure 3**

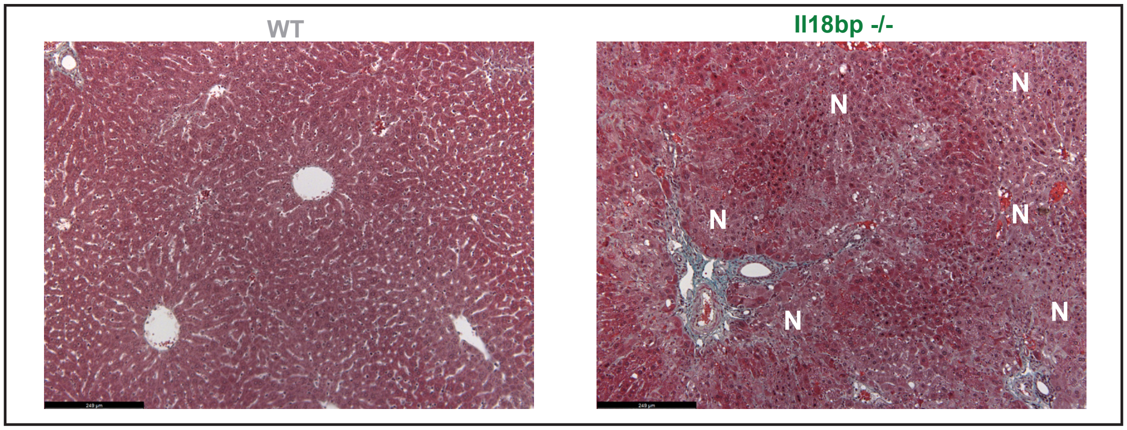


**Supplemental Figure 4**

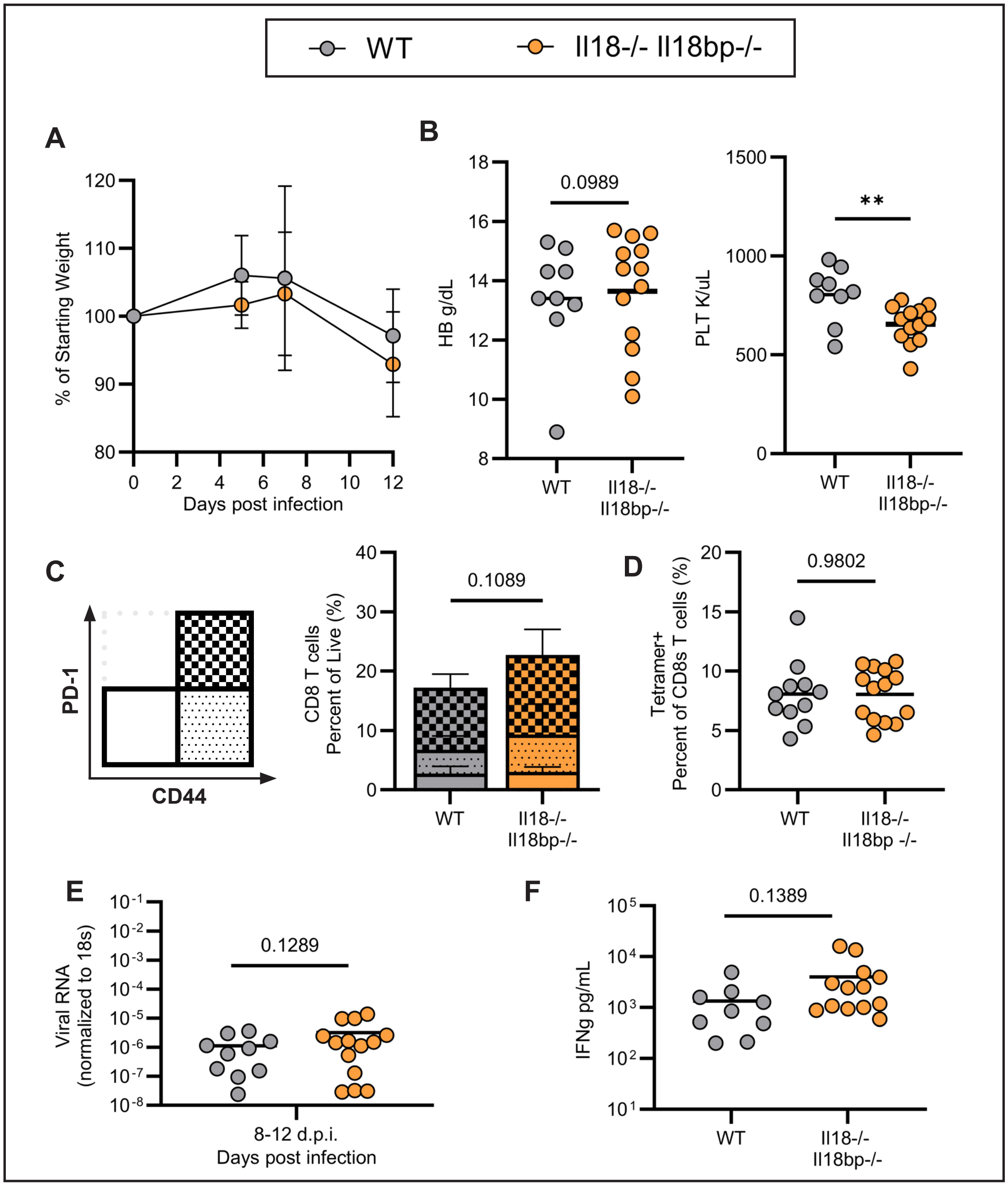


Supplemental Figure 5
