## Supplemental Methods for "Multimodal induction of fulminant HLH by IL-18 includes virus-specific NK immunodeficiency"

**Complete Medium**: RPMI 1640 with 2.05mM L-Glutamine (Cytiva HyClone) supplemented with 10mM HEPES (Gibco), 10% FBS (GenClone), 100U/mL Penn Strep (Gibco), 10mM Non-essential Amino Acids (Gibco), 0.05mM of B2-mercaptoethanol (Gibco), and 1mM Sodium Pyruvate (Gibco).

**Custom gene sets: NK IL-18 response:** Alongside the NK cell ex vivo cultures performed in *Sciume et al*^47^, sorted primary murine splenic NK cells were separately cultured with [IL-18 alone] (50ng/mL, R&D) and [IL-18 + IL-12] (IL-12 10ng/mL, R&D) for 6 hours before preparation of bulk RNAseq libraries. We combined samples from IL-18-exposed conditions ([IL-18 alone] and [IL-18 + IL-12]) and control conditions ([media alone], [IL-12 alone], and [IL-2+IL12]), and generated “Up with 18” and “Down with 18” gene sets by comparing these two groups and extracting genes with mean TPM>5 (in either group) and False Discovery Rate p-value < 0.001 (CLC genomics software as above, **Supplemental File 2**). GSEA was performed as above using these custom gene sets. **Immature NK/ILC**: We also generated custom gene sets of genes Up and Downregulated in immature vs mature NK-like cells by comparing relevant bulk RNAseq datasets from GSE77695^48^: “Immature Liver NK”=BM_iNK vs. Lv_NK, “Immature Spleen NK”= BM_iNK vs. Sp_NK, and “Immature Liver ILC1” BM_NKp vs. Lv_ILC1. We extracted genes with p < 0.01 by uncorrected t-test and FC as identified in (**Supplemental File 3**).

**Antibodies used for flow cytometry**

| **Marker** | **Fluorophore** | **Clone** | **Manufacturer** | **Catalog Number** |
| --- | --- | --- | --- | --- |
| B220/ CD45R | BV605 | RA3-6B2 | Biolegend | 103243 |
| B8R H2-Kb (sequence: TSYKFESV) | APC |  | NIH Tetramer Core |  |
| CD11b | BV711 | M1/70 | Biolegend | 101241 |
| CD11b/ MAC-1 | APC-Cy7 | M1/70 | Biolegend | 101225 |
| CD127/ IL-7r | BV605 | A7R34 | Biolegend | 135025 |
| CD127/ IL-7r | PE-Cy7 | SB/199 | Biolegend | 121120 |
| CD178 (Fas Ligand) | APC-eFluor™ 780 | MFL3 | eBio | 47-5911-82 |
| CD19 | APC | 1D3/CD19 | Biolegend | 152409 |
| CD19 | PE/DAZZLE594 | 6D5 | Biolegend | 115553 |
| CD253 (TRAIL) | Biotin | N2B2 | Biolegend | 109303 |
| CD27 | Pacific Blue | LG.3A10 | Biolegend | 124217 |
| CD279 (PD-1) | PerCP-e710 | RMP1-30 | eBio | 46-9981-80 |
| CD279/PD-1 | BV421 | RMP1-30 | Biolegend | 109121 |
| CD279/PD-1 | PerCP-Cy5.5 | RMP1-30 | Biolegend | 109119 |
| CD279/PD-1 | PE-Dazzle | RMP1-30 | Biolegend | 109115 |
| CD314 (NKG2D) | PE | CX5 | BioLegend | 130207 |
| CD335/NKp46 | APC | 29A1.4 | Biolegend | 137627 |
| CD335/NKp46 | AF647 | 29A1.4 | BD | 560755 |
| CD335/NKp46 | BV605 | 29A1.4 | Biolegend | 137619 |
| CD39 | BV605 | Y23-1185 | BD | 567266 |
| CD39 | PE-Cy7 | Duha59 | Biolegend | 143805 |
| CD4 | BUV737 | RM4-5 | BD | 612844 |
| CD4 | BV785 | GK1.5 | BioLegend | 100453 |
| CD4 | FITC | RM4-5 | Biolegend | 100509 |
| CD4 | PerCP-Cy5.5 | RM4-5 | BioLegend | 116012 |
| CD4 | APC-Cy7 | RM4-5 | Tonbo | 25-0042 |
| CD44 | BV650 | IM7 | Biolegend | 103049 |
| CD44 | PerCP-Cy5.5 | IM7 | BD | 560570 |
| CD44 | AF700 | IM7 | bd | 560567 |
| CD49b | APC-Cy7 | DX5 | Biolegend | 108920 |
| CD62L | BV711 | MEL-14 | Biolegend | 104445 |
| CD8a | Pacific Blue | 53-6.7 | Biolegend | 100728 |
| CD8a | PerCP-Cy5.5 | 53-6.7 | eBio | 45-0081-82 |
| CD8a | AF700 | 53-6.7 | BD | 557959 |
| CD8a | APC-Cy7 | 53-6.7 | BD | 557654 |
| CD8a | APC-eFluor™ 780 | 53-6.7 | eBio | 47-0081-80 |
| CD8b | APC-Cy7 | YTS156.7.7 | Biolegend | 126619 |
| CD8b/Ly3 | PE-Dazzle | YTS 156.7.7 | BioLegend | 126621 |
| FC Block |  | 93 | Biolegend | 101320 |
| IL-18Ra (CD218a) | APC | P3TUNYA | eBio | 17-5183-82 |
| IL-18Ra (CD218a) | PE-Cy7 | P3TUNYA | Thermo Fisher | 25-5183-82 |
| IL-18Ra (CD218a) | PE | P3TUNYA | eBio | 2460173 |
| KLRG1 | BV785 | 2F1/KLRG1 | Biolegend | 138429 |
| KLRG1 | PE-Cy7 | 2F1/KLRG1 | Biolegend | 138416 |
| KLRG1 | PE | 2F1 | eBio | 12-5893-80 |
| LAG-3 | BV650 | C9B7W | Biolegend | 125257 |
| Fixable LIVE/DEAD | Aqua |  | Invitrogen | L34957 |
| Ly6C | AF700 | HK1.4 | Biolegend | 128024 |
| NK1.1 | BV421 | pk136 | Biolegend | 108731 |
| NK1.1 | FITC | PK136 | Biolegend | 108705 |
| NK1.1 | PerCP-Cy5.5 | PK136 | Biolegend | 108727 |
| NK1.1 | APC-eFluor™ 780 | PK136 | eBio | 47-5941-82 |
| NK1.1 | APC | PK136 | Tonbo | 20-5941 |
| NK1.1 | PE-Cy7 | PK136 | eBio | 25-5941-82 |
| NK1.1 | PE-Dazzle | PK136 | Biolegend | 108747 |
| Streptavidin | BV421 |  | Biolegend | 405225 |
| Streptavidin | BV605 |  | Biolegend | 405229 |
| TCRb | Pacific Blue | H57-597 | Biolegend | 109225 |
| TCRb | BUV737 | H57-597 | BD | 612821 |
| TCRb | FITC | H57-597 | Tonbo | 35-5961 |
| TCRb | AF700 | H57-597 | Biolegend | 109224 |
| TCRb | APC | H57-597 | Biolegend | 109212 |
| TCRb | PE-Cy7 | H57-597 | Tonbo | 60-5961 |
| TIGIT | BV421 | 1G9 | Biolegend | 142111 |
